# The temporal dynamics of lncRNA *Firre*-mediated epigenetic and transcriptional regulation

**DOI:** 10.1101/2022.05.15.492001

**Authors:** Christian Much, Erika L. Lasda, Isabela T. Pereira, Tenaya K. Vallery, Daniel Ramirez, Jordan P. Lewandowski, Robin D. Dowell, Michael J. Smallegan, John L. Rinn

**Affiliations:** BioFrontiers Institute, University of Colorado Boulder, Boulder, CO 80303, USA; Department of Stem Cell and Regenerative Biology, Harvard University, Cambridge, MA 02138, USA; Department of Molecular, Cellular and Developmental Biology, University of Colorado Boulder, Boulder, CO 80302, USA; Department of Biochemistry, University of Colorado Boulder, Boulder, CO 80303, USA

## Abstract

Numerous studies have now demonstrated that lncRNAs can influence gene expression programs leading to cell and organismal phenotypes. Typically, lncRNA perturbations and concomitant changes in gene expression are measured on the timescale of many hours to days. Thus, we currently lack a temporally grounded understanding of the primary, secondary, and tertiary relationships of lncRNA-mediated transcriptional and epigenetic regulation – despite being a prerequisite to elucidating lncRNA mechanisms. To begin to address when and where a lncRNA regulates gene expression, we genetically engineered cell lines to temporally induce the lncRNA *Firre*. Using this approach, we were able to monitor lncRNA transcriptional regulatory events from 15 min to four days. We observed that upon induction, *Firre* RNA regulates epigenetic and transcriptional states, *in trans*, within 30 min. These early regulatory events result in much larger transcriptional changes after twelve hours, well before current studies monitor lncRNA regulation. Moreover, *Firre*-mediated gene expression changes are epigenetically remembered for days. Overall, this study suggests that lncRNAs can rapidly regulate gene expression by establishing persistent epigenetic and transcriptional states.

## Introduction

To date, dozens of lncRNAs have been found to contribute to a variety of phenotypes *in vivo*^1^. In contrast, the gene targets directly regulated by lncRNAs resulting in these phenotypes are far less understood. Most loss- or gain-of-function studies of lncRNAs measure the transcriptome at a single time point on the order of multiple hours to days or at homeostasis (e.g., knockout models). At such timescales, it is typically observed that lncRNAs regulate hundreds to thousands of genes. Thus, the earliest or primary effects of the lncRNA regulation are obscured by layers of gene-regulatory cascades and compensation mechanisms.

This has led to a variety of mechanistic models that would require timescales of many hours to days^1–19^. Yet, if a lncRNA regulates gene expression rapidly within minutes, it will limit possible mechanisms and may rule out many of the existing models of lncRNA-based regulation. Thus, determining the earliest, or most direct, regulatory targets of lncRNAs will provide a basis for resolving temporally grounded mechanistic models. Currently, we lack an understanding of the primary, secondary, and further downstream regulatory events mediated by lncRNAs in order to identify the most primary regulatory sites.

Here we set out to finely map the temporal regulatory steps mediated by a lncRNA, in *trans*, from 15 min to four days. We chose the lncRNA *Firre* (Functional intergenic repeating RNA element) based on the following properties: (i) *Firre* is a *trans*-acting RNA that does not regulate gene expression *in cis*; (ii) gain- and loss-of-*Firre* RNA function *in vivo* phenotypes range from lethality under lipopolysaccharide exposure to defects in hematopoiesis; (iii) *Firre* is genetically associated with human diseases^20–24^. Although many studies have attempted to understand the mechanism of *Firre*, these efforts have resulted in several conflicting models – a conundrum that the knowledge of *Firre*’s primary regulatory sites would help to resolve^23,25–27^.

To this end, we engineered an inducible *Firre* mouse embryonic stem cell (mESC) system in a *Firre* knockout and wild-type (WT) background. We monitored gene expression across 16 time points on the order of minutes, hours, and days after induction of *Firre*. Combining RNA-seq, ATAC-seq, and PRO-seq across relevant time windows, we found that *Firre* can regulate transcription in a matter of minutes. Moreover, we observed changes in epigenetic state and nascent transcription within 30 min and mature target gene product within two hours of *Firre* induction. At longer timescales, we detected numerous gene expression changes that are likely secondary or downstream modes of gene regulation. Together, our results suggest that *Firre* functions via an RNA-based mechanism that can transduce a signal within minutes to change epigenetic state and activate gene expression that then persists for days.

## Results

### Genetically engineered mESC lines for temporal control of *Firre*

To understand the temporal and sequential regulatory role of the *Firre* lncRNA, we need to be able to control the expression of *Firre* and measure the dynamics of gene expression changes before the cell returns to homeostasis. While inducible transgenes have the advantage of determining temporally grounded regulatory events, there are several key caveats to consider when using these transgenic systems. This includes the amount of induction over physiological levels and the off-target effects of doxycycline (dox).

To control for varying levels of *Firre* induction, we generated two dox-inducible *Firre* transgene lines (Fig. 1A, Supplemental Fig. S1A). For both lines, we chose the *Firre* isoform (1.49) that was previously shown to have functional roles *in vivo*^20^. One mESC line contains an inducible *Firre* transgene in a *Firre* knockout background that we term *Firre^RESCUE^* (Fig. 1A). The other one has the *Firre* transgene in a WT background, termed *Firre^OE^*. To control for effects of dox itself in the *Firre^RESCUE^* line, we generated an mESC line termed *Firre^KO-CTL^* that lacks endogenous *Firre* and contains the *Firre* transgene but does not express the reverse tetracycline-controlled transactivator (rtTA) element (Fig. 1A, Supplemental Fig. S1B). To control for dox in the *Firre^OE^*line, we used WT cells.

**Figure 1:**
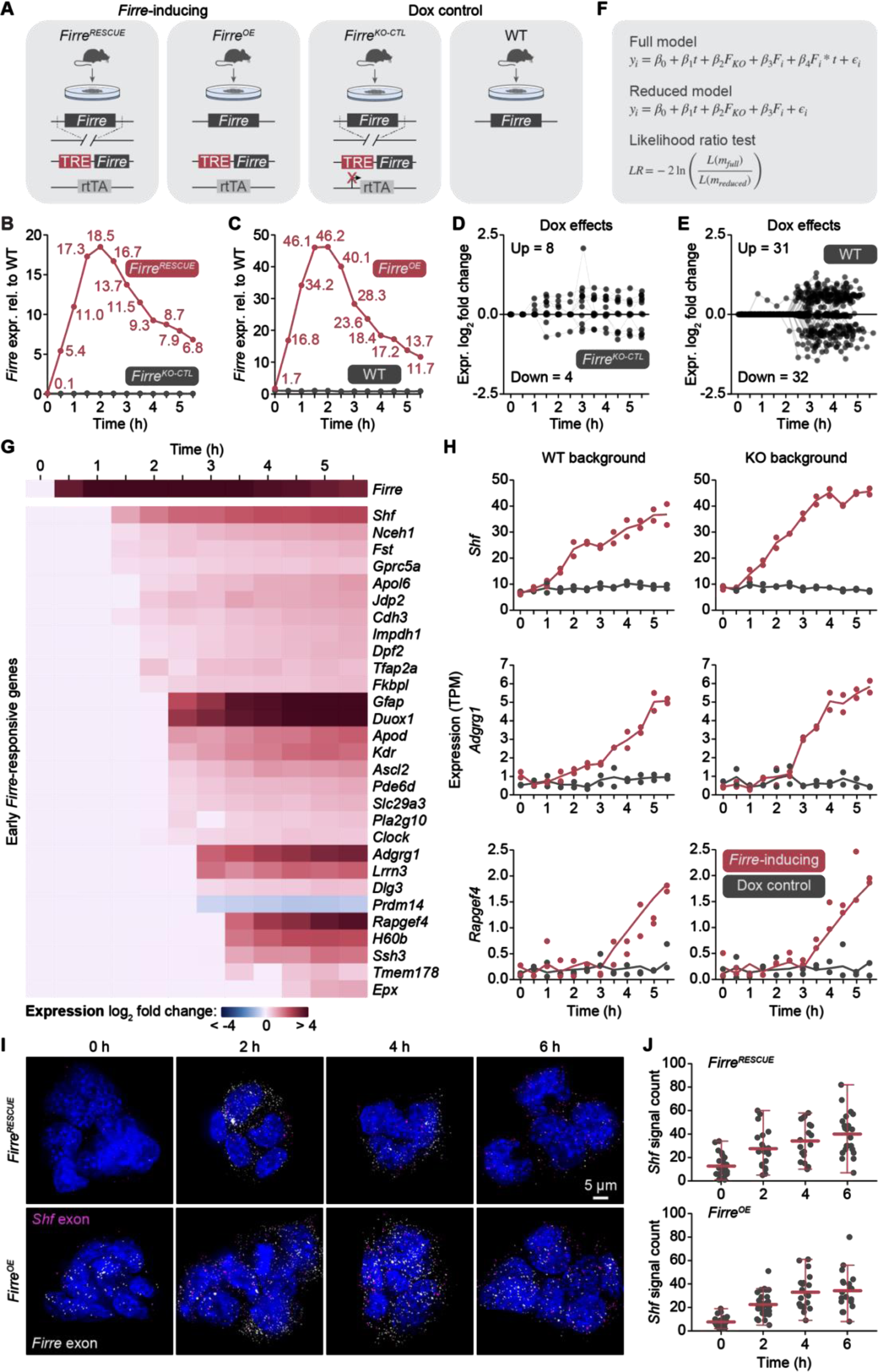
Temporal gene regulation in *Firre* transgene mESC lines. **(A)** Schematic showing the mouse blastocyst-derived mESC lines used in this study. The *Firre^RESCUE^* and *Firre^OE^* cell lines contain a dox-inducible *Firre* transgene in a *Firre* KO and WT background, respectively. The *Firre^KO-CTL^* and WT cell lines do not induce *Firre* expression and serve as dox controls. **(B, C)** *Firre* expression levels in the *Firre^RESCUE^* (B) and *Firre^OE^* (C) line (red) as well as the *Firre^KO-CTL^* (B) and WT (C) line (black) across time after the addition of dox as measured by RNA-seq. Numbers indicate fold expression changes over *Firre* WT levels at each time point. **(D, E)** Expression log_2_ fold change of significantly differentially expressed genes that are affected by dox treatment in the *Firre^KO-CTL^* (D) and WT (E) control cell line. Fold changes are relative to the zero-time point.**(F)** Linear model to derive genes that are significantly differentially regulated upon *Firre* transgene induction and not due to dox treatment (likelihood ratio test). **(G)** Heatmap of gene expression changes of early *Firre*-responsive genes over time as determined by the linear model in (F). **(H)** Gene expression changes of representative genes due to *Firre* in *Firre*-inducing (red) but not in dox control (black) cells of *Firre* WT and KO background. **(I)** Maximum intensity projections of smRNA FISH images for *Shf* exon (magenta) and *Firre* exon signals (white) in *Firre^RESCUE^*and *Firre^OE^* mESC colonies after 0, 2, 4, and 6 hours of dox treatment. Nuclei are stained with Hoechst (blue), *Firre* exon in white, and *Shf* exon in magenta. Scale bar is 5 μm. (**J**) Quantification of *Shf* exon FISH signals in *Firre^RESCUE^* and *Firre^OE^*mESCs. Data are mean and non-outlier range of quantified cells.

To validate *Firre* expression upon induction in our cell lines, we performed RT-qPCR using *Firre* probes detecting both endogenous and transgene *Firre* after zero and twelve hours of dox treatment. In the *Firre^OE^* line, *Firre* expression was induced to about 2.5-fold above WT levels, while in the *Firre^RESCUE^*line, *Firre* expression reached WT levels after twelve hours. Importantly, we did not observe any induction of *Firre* expression in the WT and *Firre^KO-CTL^* line (Supplemental Fig. S1C). Of note, we confirmed that neither the induction of *Firre* nor dox administration impacted mESC growth or morphology (Supplemental Fig. S1D). Collectively, we generated cell lines to temporally track varying levels of *Firre* expression and identify genes that are regulated by *Firre* and not dox.

### Early *Firre*-mediated gene regulation

To test the temporal dynamics of each of these inducible lines, we collected RNA every 30 min until 5.5 hours post induction of *Firre* by addition of dox and performed RNA-seq with two replicates for each time point. In both the *Firre^RESCUE^* and *Firre^OE^* lines, we detected induction of *Firre* expression already within 30 min and peak expression at around two hours (Fig. 1B-C). As expected, we observed varying levels of *Firre* at different time points in the *Firre^RESCUE^* and *Firre^OE^* lines. This property will further allow us to identify genes regulated by *Firre* across a spectrum of induction levels.

We next investigated which genes were regulated by dox itself and not due to the induction of the *Firre* lncRNA. In both *Firre^KO-CTL^* and WT control models, we detected very few genes that were regulated by dox alone (*Firre^KO-CTL^*= 12 genes, WT = 63 genes; Fig. 1D-E). Moreover, we observed that most dox-based artifacts occurred after approximately 2.5 hours in both control models. Thus, any regulatory events observed prior would be due to the *Firre* lncRNA and not due to artifacts of the transgene system.

To generate a high-confidence list of *Firre*-mediated transcriptional regulatory events, we used a likelihood ratio test (LRT) between two regression models (Fig. 1F, see methods). The first regression model contains parameters comparing genetic background (*Firre^RESCUE^* and *Firre^OE^* models), dox control lines, and time. The second regression model lacks the dox control lines and models genetic background and time. Thus, this conservative approach requires target genes to change significantly in time, in both WT and KO background, and not due to dox. Using our conservative LRT statistical approach, we identified *Firre* as a positive control as well as 29 early *Firre*-responsive genes concordantly regulated in both *Firre* WT and KO backgrounds (P < 0.05, |fold change| > 1.5; Fig. 1G-H). Notably, all but one (*Prdm14*) of these genes were upregulated, starting to increase in expression in 1.5-4.5 hours (Fig. 1G-H). Given the short gene list, gene ontology (GO) analysis determined no meaningful common pathways or processes among the 29 early *Firre*-regulatory targets (Supplemental Fig. S1E). Collectively, we identified a cohort of genes that are the earliest and most robust responders to the induction of *Firre* RNA at different levels in WT and KO backgrounds.

While our control cell lines account for dox, they do not control for the role of the rtTA protein after dox activation. To ensure that the early *Firre* target genes are not artifacts of rtTA activation by dox, we analyzed gene expression data in mESCs with a dox-inducible Nanog transgene generated by Heurtier et al.^28^. We found that none of the early *Firre*-responsive genes changed significantly upon activation of rtTA by dox treatment. This result indicates that the early *Firre* targets were not induced as an artefact of rtTA activation but rather due to the *Firre* RNA (Supplemental Fig. S1F).

In addition to the dox controls above, we further wanted to validate the temporal regulatory events by single molecule RNA FISH (smRNA FISH). We chose *Shf* and designed a complementary exon probe set to monitor its RNA abundance and localization across time. We tested 0, 2, 4, and 6-hour time points after *Firre* induction in *Firre^RESCUE^* and *Firre^OE^* lines. In both cell lines, we observed a continuous increase in the expression of *Shf* after the induction of *Firre*, independently validating our RNA-seq results with an orthogonal methodology (Fig. 1I-J). Collectively, these data point to 29 high-confidence genes that are rapidly and robustly changing upon exposure to varying levels of *Firre* RNA across time and not due to dox or rtTA. Thus, we have likely identified the primary regulatory sites of the *Firre* lncRNA that occur at timescales hitherto unstudied.

### Temporal dynamics of epigenetic regulation by *Firre*

Based on our finding that *Firre* regulates primary target sites within a few hours, we further hypothesized that changes to epigenetic states may occur prior to these transcriptional changes. To test this hypothesis, we performed ATAC-seq every 30 min for 2.5 hours across the four different mESC lines. Importantly, we found no ATAC peaks significantly changing in time due to dox (P < 0.05) in this early time frame. Testing for differential accessibility by comparing all conditions in which *Firre* was induced (time point > 0 in *Firre^RESCUE^* and *Firre^OE^* lines) to conditions in which the *Firre* transgene was not induced (WT and *Firre^KO-CTL^*lines; zero-time point in *Firre^RESCUE^* and *Firre^OE^* lines), we identified a total of 55 ATAC peaks that changed (P < 0.05) in response to *Firre*’s induction (Fig. 2A, Supplemental Fig. S2A). Of these 55 peaks, 51 (93%) were associated with an increase in chromatin accessibility over the time course (Fig. 2A, Supplemental Fig. S2A), which is consistent with the observed subsequent transcriptional activation.

**Figure 2:**
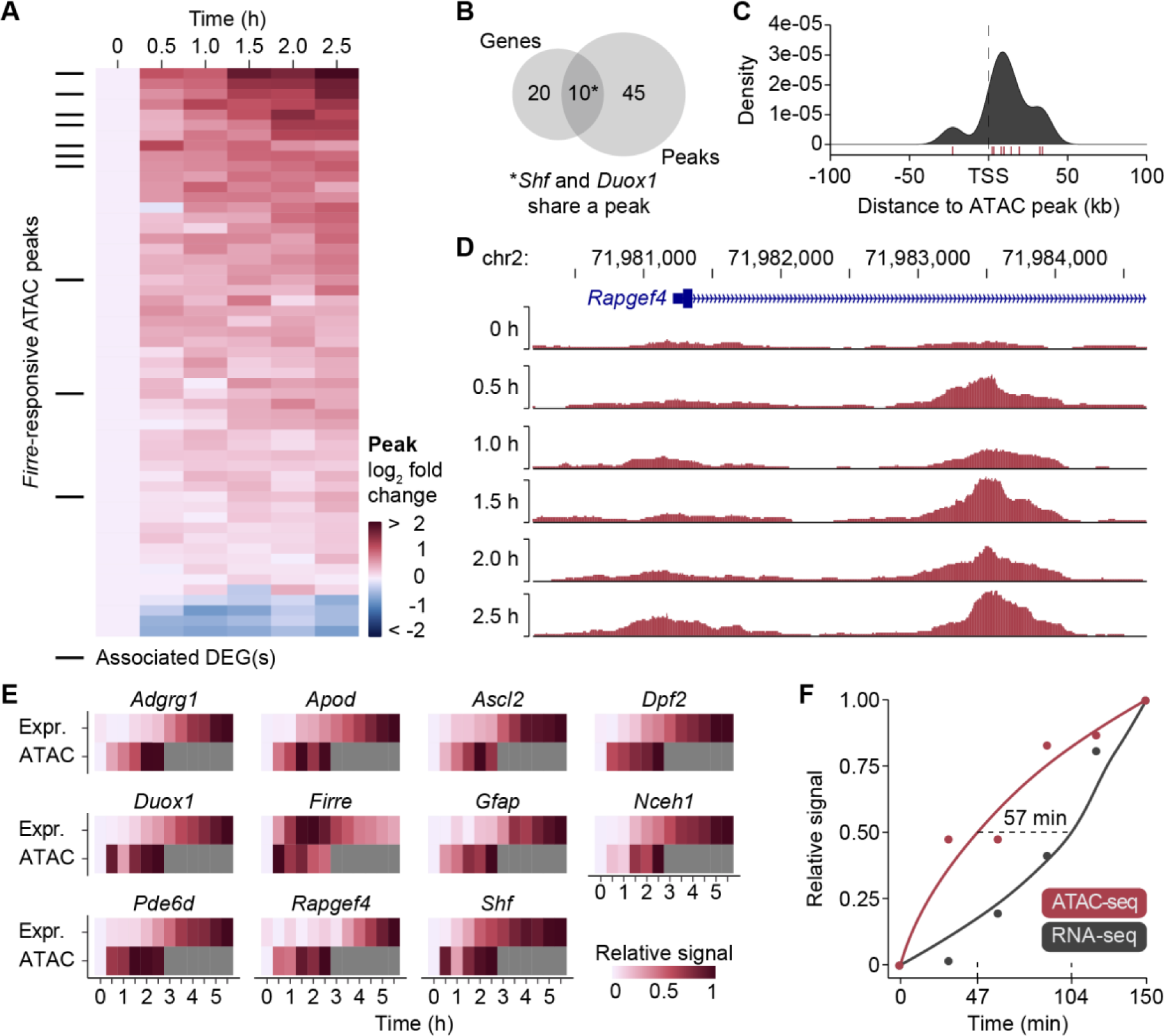
Changes in chromatin accessibility upon induction of *Firre*. **(A)** Heatmap showing chromatin accessibility log_2_ fold changes across time. Increased chromatin accessibility relative to the zero-time point is depicted in shades of red, decreased chromatin accessibility in blue. Black tick marks indicate differentially expressed genes. **(B)** Venn diagram showing the overlap of early *Firre*-responsive genes and *Firre*-mediated chromatin accessibility sites. **(C)** TSS metaplot of the density of ATAC-seq peak centers relative to *Firre*-responsive gene TSSs. Red tick marks indicate the peak centers. **(D)** Genome browser tracks of chromatin accessibility changes across time within *Rapgef4* in *Firre^RESCUE^*cells. **(E)** Heatmap of chromatin accessibility and gene expression changes at each time point after induction of *Firre.* Time points where chromatin accessibility was not measured are indicated in grey. **(F)** Plot of chromatin accessibility (red) and gene expression (black) signal relative to the maximum measured over time. The dashed line represents the time difference in half-maximum values.

Next, we wanted to determine how many of the differentially accessible ATAC peaks were in proximity to genes changing in expression upon *Firre* induction. We identified ten differentially accessible ATAC peaks within 50 kb of the transcription start site (TSS) of early *Firre*-responsive genes and *Firre* itself (Fig. 2B). One of these peaks was in proximity to two neighboring genes *Duox1* and *Shf*, which both increased in expression. Analysis of the ATAC-seq peak center location relative to the TSS for the ten genes (excluding *Firre*) revealed that all but one ATAC peaks were downstream of the TSS (Fig. 2C). The gene *Rapgef4*, for example, rapidly gained ATAC peak signals within its first intron at a proximal enhancer element (Fig. 2D, Supplemental Fig. S2B). Together, these results suggest that the *Firre* RNA is sufficient to alter chromatin accessibility within 30 min.

We next wanted to define the timing of chromatin accessibility and gene expression changes upon *Firre* induction. To this end, we compared the temporal dynamics of ATAC- and RNA-seq changes side by side. We found that for all eleven genes (including *Firre*), chromatin accessibility increased already within 30 min and in all cases preceded gene expression changes (Fig. 2E). Importantly, within this time window, *Firre* was expressed at or near WT levels in both *Firre*-inducing cell lines. To formalize the observed temporal dynamics, we created a meta-profile of ATAC- and RNA-seq signal changes for these genes (Fig. 2F). We found that, on average, ATAC peaks and RNA abundance reached half-maximal at 47 min and 104 min, respectively. Together, these data provide evidence that *Firre* rapidly increases chromatin accessibility which in turn leads to an upregulation of gene expression.

### *Firre* regulates RNA transcription within minutes

Our results above demonstrate that the *Firre* RNA can induce changes in chromatin within 30 min, suggesting that transcriptional changes might be occurring on short timescales as well. As poly(A) RNA-seq is only detecting mature transcripts, its temporal resolution is limited by pre-mRNA processing time. Therefore, to identify transcriptional regulatory events at or prior to 30 min, we performed PRO-seq which measures nascent transcription with a temporal resolution on the order of minutes^29^. Importantly, at these time points, *Firre* is induced to near-WT levels in the *Firre^RESCUE^* line. Thus, any regulatory events observed are likely not a consequence of *Firre* overexpression, but rather due to *Firre* RNA expressed at its physiological levels.

To examine the earliest and most direct regulatory effects of *Firre*, we performed PRO-seq in the *Firre^RESCUE^*model at 0, 15, and 30 min post addition of dox in biological triplicates. We first used Tfit^30^ to determine the number of bidirectional regions indicative of transcriptional activity (Fig. 3A). This resulted in a total of 29,842 sites of bidirectional transcription. Most of these sites of RNA polymerase activity remained unchanged, including the *Ep300* promoter (Fig. 3B). However, at genes differentially expressed at the mature RNA level, such as *Shf* and *Duox1*, we interestingly observed changes in nascent transcription within 15 min (Fig. 3B).

**Figure 3:**
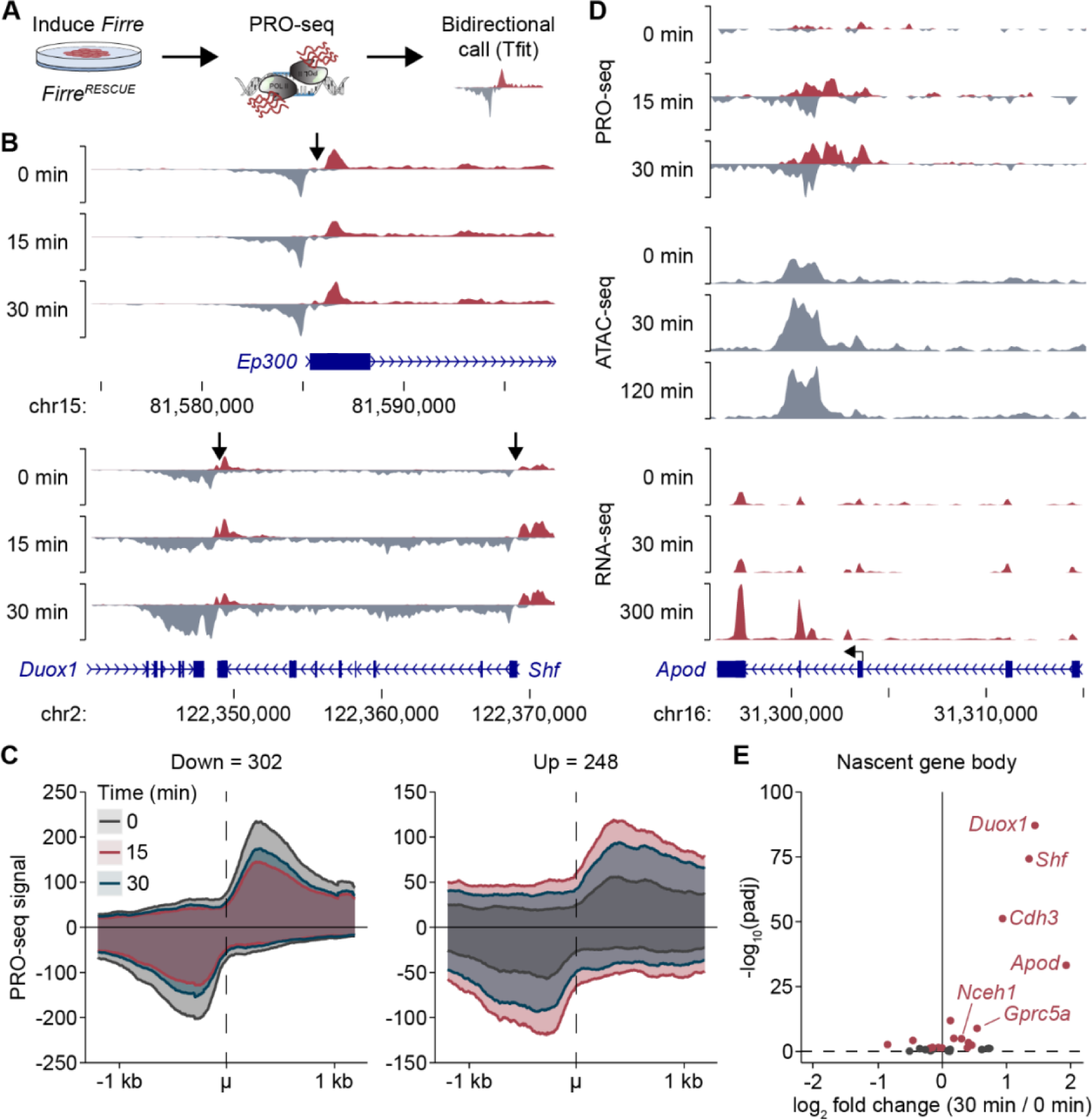
Rapid nascent transcriptional changes upon *Firre* induction. **(A)** Schematic of PRO-seq experimental setup. **(B)** Genome browser tracks of bidirectional transcription of genes that were (*Shf*, *Duox1*) and were not (*Ep300*) differentially expressed upon *Firre* induction. Red and grey areas indicate reads on the plus and minus strand, respectively. Arrows denote the bidirectional center. **(C)** Metaplot of PRO-seq signal at the differentially expressed sites of bidirectional transcription (padj < 0.05, n = 550) around the bidirectional center (μ) at the 0-, 15-, and 30-min time points. **(D)** Genome browser tracks showing the regulatory events at the *Apod* locus upon *Firre* induction. The arrow indicates an internal TSS. **(E)** Volcano plot of the log_2_ fold changes of nascent transcription over the gene body at 30 vs 0 min. Genes with padj < 0.05 are indicated in red.

To determine differentially expressed bidirectional sites globally, we performed an LRT by comparing a full model with the time component to a reduced model representing the mean expression, resulting in 550 differentially expressed bidirectional regions (padj < 0.05; |fold change| >= 1.5). In contrast with the observed changes in mature RNA being predominantly gene activation, sites of bidirectional nascent transcription were up- and downregulated (upregulated = 248, downregulated = 302; Fig. 3C, Supplemental Fig. S3A). Most bidirectional sites of nascent transcription responded transiently at 15 min, followed by a return toward pre-induction levels by 30 min. However, 41 of the 550 differentially expressed bidirectionals continued a monotonic trend. Of those, *Apod*, *Shf*, and *Duox1* had monotonically increasing bidirectional signals co-located within increasing ATAC-seq peaks (Fig. 3D). Correspondingly, these regions exhibited some of the largest expression fold changes in mature transcripts. Thus, we can now place *Firre*- regulatory events in temporal sequence of primary chromatin accessibility targets that lead to the subsequent mRNA changes shortly after.

To assess the concordance between nascent transcription and RNA-seq expression changes, we calculated the differential expression for nascent transcripts along the gene body. We detected 18 of the 29 early *Firre* responders as significantly differentially expressed at the nascent transcript level (padj < 0.05, Fig 3E). The five genes with the earliest detected changes at 1.5 hours in RNA-seq (*Shf*, *Nceh1*, *Fst*, *Gprc5a*, and *Cdh3*) all had increasing nascent transcription along the gene body by 30 min, and four of those five genes were significantly upregulated one to two hours later (padj < 0.05, Fig 3E, Supplemental Fig. S3B). In summary, we identified many sites that rapidly and transiently respond to *Firre*, with only some resulting in persistent nascent and mature expression changes. This underscores the importance of multiple genomic scale approaches to home in on bona fide lncRNA gene targets that exhibit persistent opening of chromatin, induction of nascent transcription and robust activation of the target gene’s mature transcript.

### *Firre*-mediated transcriptional regulation persists for days

While the goal of this study is to identify robust, reproducible, and primary regulatory sites of *Firre*, we also wanted to determine how long these regulatory changes last. To this end, we performed an RNA-seq time course for 0, 12, 24, 48, and 96 hours in biological triplicate for all four cell lines at each time point. We used a similar LRT that requires genes to significantly change across time in both *Firre^RESCUE^* and *Firre^OE^* lines and neither in *Firre^KO-CTL^* nor WT cells treated with dox. We observed that in both the *Firre^RESCUE^* and *Firre^OE^* lines, *Firre* was induced highest at twelve hours and then remained at much lower levels up to 96 hours (Fig. 4A-B).

**Figure 4:**
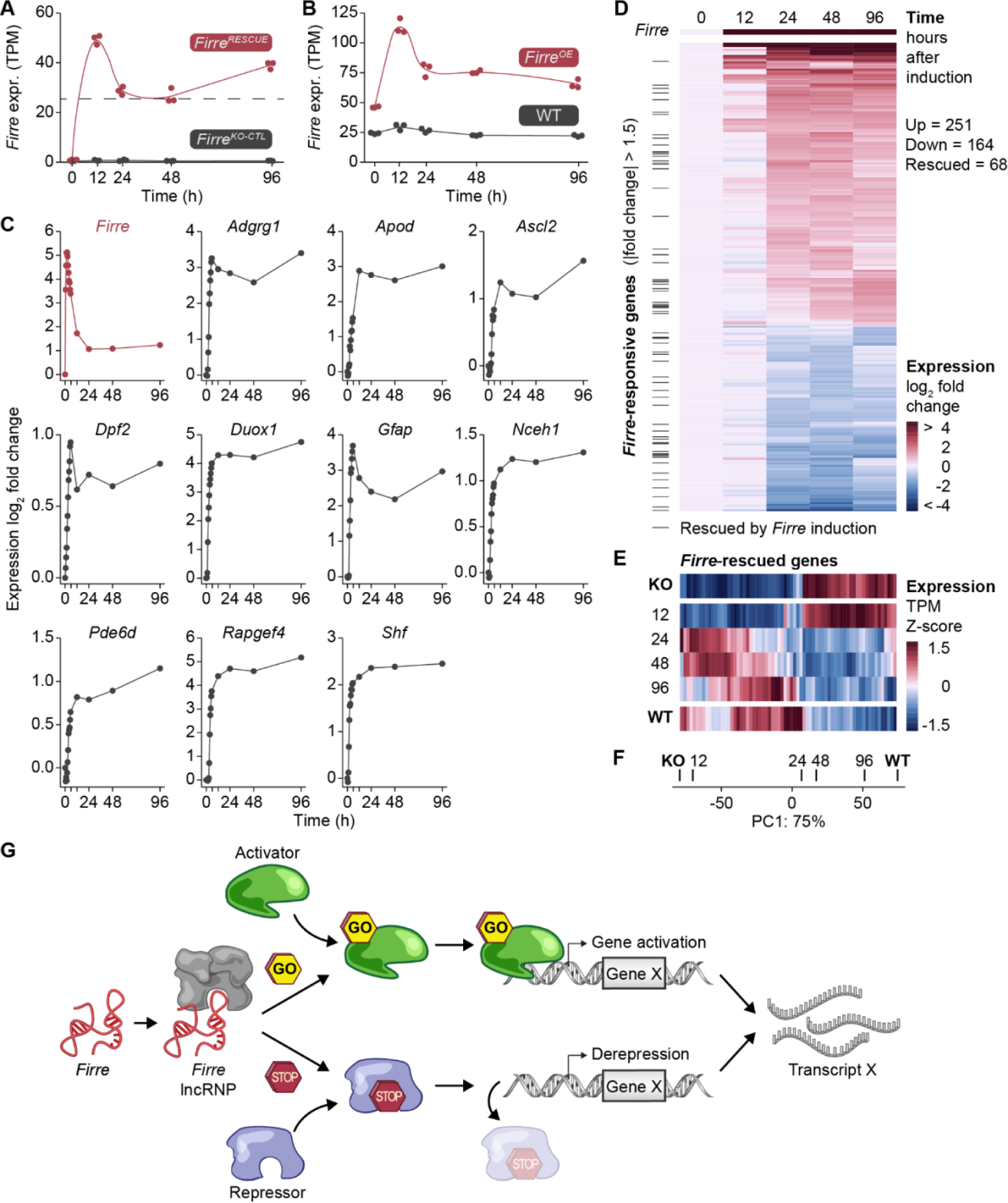
Persistence of *Firre*-mediated transcriptional regulation over time. **(A, B)** Expression levels of *Firre* in *Firre^RESCUE^* (red) and *Firre^KO-CTL^* mESCs (black) (A) and *Firre^OE^* (red) and WT mESCs (black) (B) across time after the addition of dox as measured by RNA-seq. The dashed line indicates *Firre* WT expression levels. **(C)** Individual gene plots showing the log_2_ expression fold change in the *Firre*-inducing cell lines compared to the zero-time point for all measured time points. **(D)** Heatmap of the expression fold changes of significantly differentially expressed genes over time upon *Firre* induction independent of dox effects. Black tick marks represent genes whose expression was rescued, i.e., that were downregulated in the knockout and upregulated upon transgene induction, or vice versa. **(E)** Heatmap of the expression of 68 rescued genes across each time point. Top and bottom heatmap represent the *Firre^KO^* and WT expression levels, respectively. The heatmaps in between represent the temporal timing of when genes are rescued. **(F)** PCA of the rescued gene expression levels in WT, *Firre^KO^* and *Firre^RESCUE^* cells at each time point. The primary component PC1 that represents 75% of the variation in the data is plotted. **(G)** Model of *Firre*-mediated gene regulation depicting *Firre* forming a lncRNP by binding to an unknown protein, which either engages an activator protein to bind to chromatin or evicts a repressor protein from chromatin, ultimately leading to the upregulation of *Firre* target genes.

Next, we wanted to determine if the early *Firre* target genes continued to be regulated on the timescale of days. We focused on the ten genes that had immediate changes in chromatin accessibility, activation of nascent transcription and subsequent mRNA activation two hours later. Strikingly, we observed that all ten genes were induced to maximum levels within twelve hours and persisted at or close to these levels for 96 hours (Fig. 4C). This suggests that the early epigenetic changes induced by *Firre* are remembered for days.

We further utilized this longer time course to identify the cascade of secondary and tertiary regulatory events that subsequently followed from the 10-29 primary sites of *Firre* regulation. We focused on the *Firre^RESCUE^* and *Firre^KO-CTL^* cell lines and applied a similar LRT that requires genes to be significantly changed across time and not dox. We observed 414 genes significantly regulated by *Firre* and not dox across time (P < 0.05; |fold change| >= 1.5; Fig. 4D, Supplemental Fig. S4A-B). Thus, we found a large expansion of the gene list from 10-29 genes at the earliest measurable time points to 414 genes at the longer time frames. These results demonstrate the importance of temporal studies as the primary early targets persist at longer time points, but would be hard to identify in the wake of over 400 genes changing by 12-96 hours – a time frame often used in lncRNA studies.

Lastly, we wanted to leverage our *Firre^RESCUE^* line to determine the extent to which the *Firre* transgene restored gene expression to WT levels – or rescued the knockout expression phenotype. Specifically, we compared the genes that were significantly differentially expressed in response to *Firre*’s induction with those that were significantly differentially expressed in uninduced *Firre^KO^*and WT cells. Of the 414 genes responding to *Firre*’s induction, a significant number of genes (102) overlapped with the 921 differentially regulated genes in the *Firre^KO^* vs WT comparison (hypergeometric P = 5.79×10^-70^, Supplemental Fig. S4C-D). We defined genes as rescued when they were downregulated in the *Firre^KO^* cells and upregulated in the *Firre^RESCUE^*cells, or vice versa, and reverted to +/-20% of WT levels. Of the 102 overlapping genes, 68 fit these rescue criteria (Supplemental Fig. S4E). We next determined the temporal dynamics of this gene expression restoration. Principal component analysis (PCA) showed an initial shift toward rescued gene expression levels between twelve and 24 hours (Fig. 4E-F). Thus, early primary targets lead to a transcriptional cascade of over 400 genes, including those that are restored to WT levels. Collectively, we present a set of experiments that tracks the regulatory effects of the *Firre* lncRNA at its target sites from the earliest measurable changes in transcriptional activity, through changes in chromatin accessibility, to increased expression of mature RNA products which persist through the latest time point at four days.

## Discussion

Knowing the primary targets of any regulatory element is critical to understand its mechanisms. Understanding when, where, and how much gene regulation occurs allows these sites to be assayed to identify the corresponding molecular componentry underlying this regulation. For example, a common first step in understanding how a transcription factor functions is to identify the primary targets and subsequent temporal dynamics of gene regulation. These primary sites can be further used to assay for a variety of functional and mechanistic studies underlying transcription factor regulation.

Similarly, lncRNAs have been implicated in regulating a variety of transcriptional programs. However, most studies measure gene-regulatory events on the timescale of many hours to days. Thus, it is difficult to discern primary regulatory sites from secondary, tertiary, and more downstream ripple events. In contrast, it is well known that transcription factors regulate their primary targets on the scale of minutes – a timescale not explored prior to this study^29,31–43^. To begin to determine the timescales at which lncRNAs regulate transcription, we combined several genome-scale approaches and employed genetically defined models and fine-grained temporal resolution from minutes to days. We reasoned that the earliest RNA-mediated regulatory events are the most direct or primary sites of lncRNA regulation.

Using our temporally controlled approach, we made several findings with implications for understanding lncRNA-mediated gene regulation. First, we were somewhat surprised at how rapidly the gene-regulatory events unfolded upon induction of the *Firre* lncRNA. We observed that robust epigenetic and transcriptional changes occurred within 30 min – a time frame hitherto unstudied in lncRNA functional studies. Second, lncRNAs have been implicated in a variety of epigenetic mechanisms, but we lack an understanding of the temporal relationships to subsequent regulatory events. Our findings suggest that *Firre* is sufficient to induce epigenetic changes, such as chromatin accessibility, prior to the activation of genes near these sites. Moreover, this further reinforces the hypothesis that lncRNAs are epigenetic regulators. Third, our study underscores the importance of understanding the temporal dynamics underlying lncRNA regulation. Specifically, in order to determine primary targets, it was paramount to measure gene-regulatory events within a few hours of *Firre* induction. The 10-29 *Firre* target genes would have otherwise been impossible to pinpoint among the 400 or more secondary and tertiary targets observed after twelve hours. Collectively, our study demonstrates that lncRNAs could act early at a few sites that, in turn, result in larger transcriptional regulatory cascades. This is akin to what is known for transcription factors.

While temporally controlled transgene systems afford numerous insights outlined above, there are several caveats that also need to be considered. Gene expression artifacts could be caused by the exposure to dox and activated rtTA. To account for these possibilities, we exposed control cell lines that did not express rtTA to dox, and referred to an existing study to rule out any gene expression changes that could be caused by dox and activated rtTA. A second important consideration is the overexpression of *Firre* above WT levels. One way of mitigating this issue is to induce *Firre* at different levels and only consider significant regulatory events that occur independent of the abundance of *Firre*. To this end, we used the *Firre^RESCUE^* and *Firre^OE^* lines that induce *Firre* at varying levels in each line and at each time point. Thus, we only considered genes to be regulated by *Firre* if they occurred at all levels (from near WT to overexpressed). A final indication that our results are not due to overexpression artifacts is the observation that the earliest regulatory events occurred between 15-30 min. Within 30 min, we observed robust changes in chromatin accessibility and induction of nascent transcription at these sites. In this time frame, *Firre* abundance was at or near WT levels in the *Firre*-rescue model, suggesting that the observed epigenetic and transcriptional regulatory events are attributable to near WT levels of *Firre*. Overall, the reported primary regulatory events occurred independent of *Firre* abundance, were consistent across time, persisted for days, and did not take place in dox control lines.

The rapidity of RNA-based regulation by *Firre* likely precludes the possibility of a new mRNA being transcribed, spliced, exported, translated, and the protein re-localized to the nucleus and, in turn, regulating epigenetic states and transcription in the timescale of 30 min. This reduces the search space toward mechanistic models that could occur in these short time frames. Regulation by *Firre* within a few minutes likely requires an existing nucleoplasmic factor that sends a rapid signal to another factor poised at the target site. Specifically, *Firre* could be activating an activator or repressing a repressor to specifically upregulate its primary target genes (Fig. 4G).

Cells can “sense” DNA damage, heat shock, hormone exposure, and even viral RNA and transduce a signal that results in gene activation or repression within minutes^29,31–41^. It is therefore an intriguing possibility that lncRNAs could also rapidly regulate transcription by acting upstream in signal transduction cascades. This would be consistent with the RIG-I RNA-sensing pathway – where RIG-I binds RNA, is activated, and triggers a specified transcriptional response of a few immediate genes^34^. Thus, lncRNAs could also serve as allosteric adapters to activate or repress signal transduction molecules in a similar manner across a diversity of signal transduction pathways (Fig. 4G). In the case of *Firre*, we could envision an RNA-protein interaction that changes the conformation of a signaling molecule to be activated and specify a targeted transcriptional response. Finally, a nuclear localized RNA, such as *Firre*^23,44^, would be an ideal molecule for these rapid and fine-tuned transcriptional responses.

In summary, our ‘multi-omic’ temporal study demonstrates that *Firre* rapidly, robustly, and reproducibly regulates transcription and epigenetic states within 30 min – resulting in the activation of a cohort of target genes. These primary regulatory sites will serve as an invaluable tool for future work on elucidating the role of RNA-based transcriptional and epigenetic regulation. Based on these high-resolution temporal insights, we are now poised with the most immediate and direct primary sites of regulation by a lncRNA. We can next leverage these sites to assay the underlying molecular modalities and mechanisms of RNA-based epigenetic and transcriptional regulation.

## Supplemental figures

**Supplemental Figure S1:**
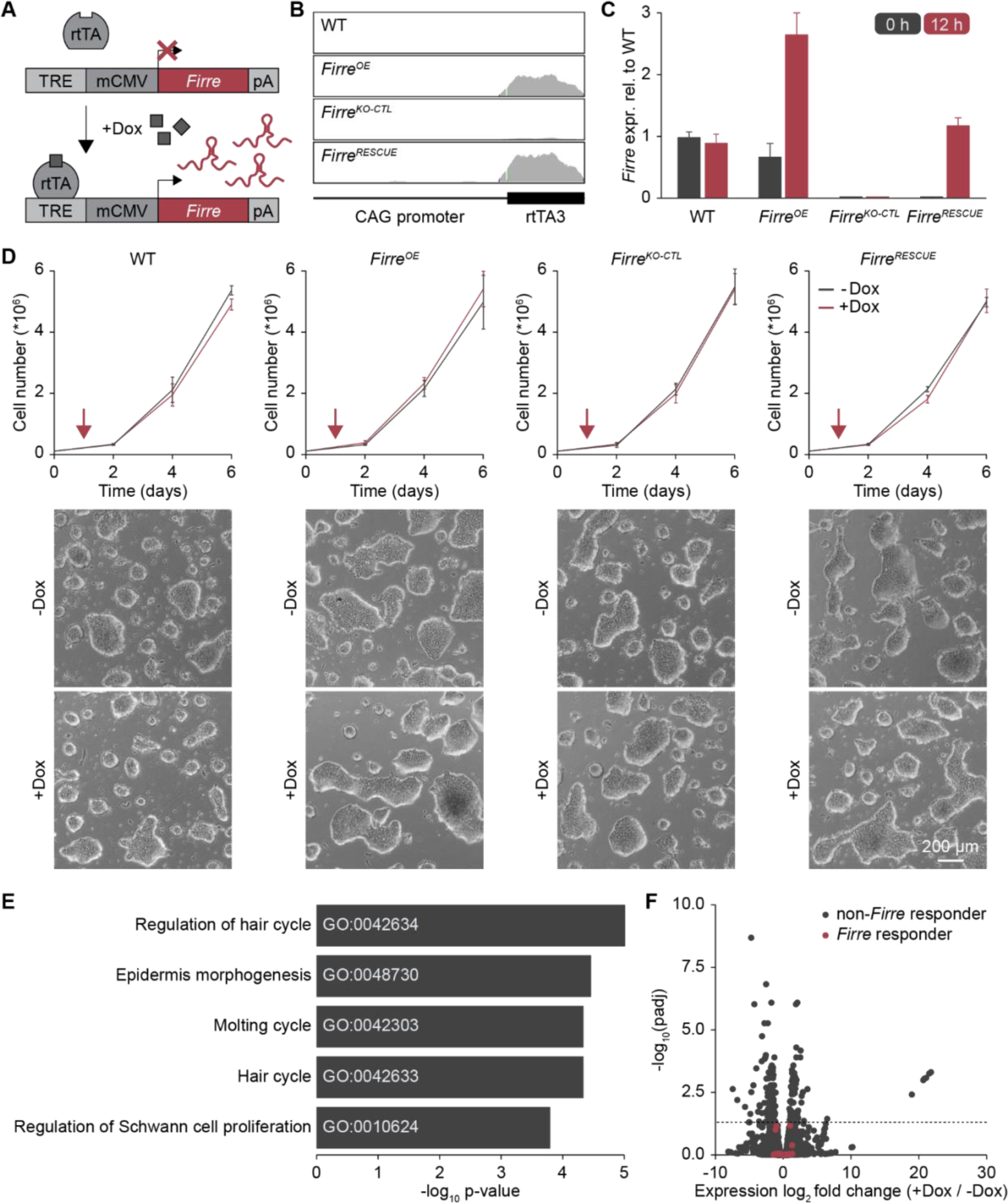
*Firre* transgene and early *Firre*-mediated gene regulation. **(A)** Diagram showing the *Firre* transgene construct and its induction by dox through the rtTA protein. **(B)** Genome browser tracks of RNA-seq reads over the rtTA3 element, which is expressed in *Firre^RESCUE^* and *Firre^OE^*cell lines, but not in the WT and *Firre^KO-CTL^* dox controls. **(C)** Relative expression of *Firre* after zero and twelve hours of dox treatment in all cell lines as determined by RT-qPCR. Expression levels have been normalized to *Firre* WT levels at zero hours. **(D)** Top: Growth curves of all cell lines treated and not treated with dox. The red arrow indicates the onset of dox treatment. The mean and standard deviation of three technical replicates are shown. Bottom: Representative images showing morphology of all cell lines treated with dox for zero and five days, respectively. Scale bar is 200 μm. **(E)** Volcano plot of the log_2_ fold expression changes in untreated and dox-treated mESCs from Heurtier et al.^28^. Early *Firre*-responsive genes are indicated in red. **(F)** GO analysis for the early *Firre*-responsive genes showing the top five most significant biological processes.

**Supplemental Figure S2:**
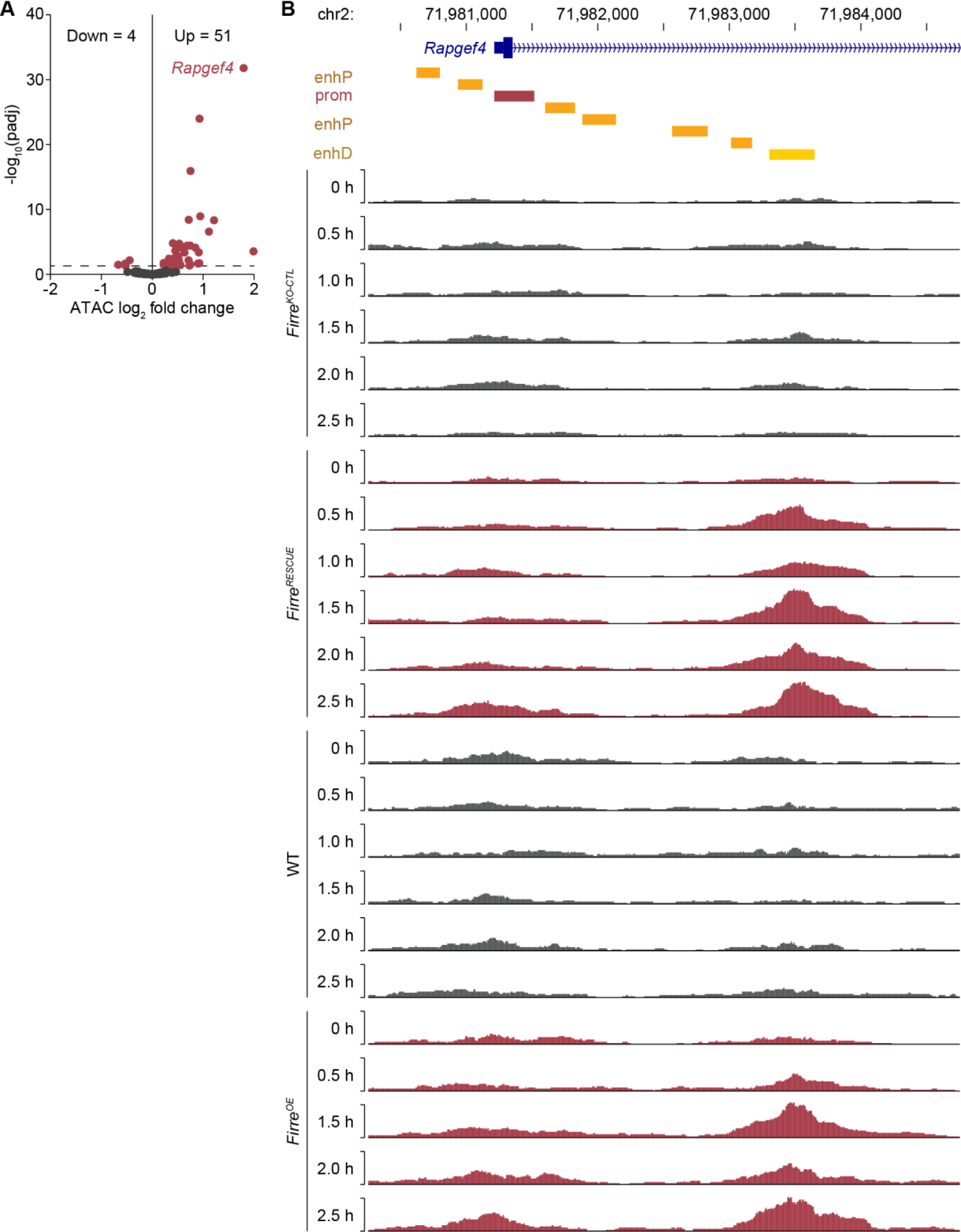
*Firre*-mediated ATAC-seq peak changes. **(A)** Volcano plot showing ATAC peak signal changes between *Firre*-inducing and dox control cells. Significantly differentially accessible peaks are plotted in red with a cutoff of padj < 0.05. **(B)** Genome browser tracks of ATAC-seq peak signal at the *Rapgef4* locus in all cell lines through time after dox administration. Predicted promoter (prom) and enhancer elements (enhP and enhD) are depicted.

**Supplemental Figure S3:**
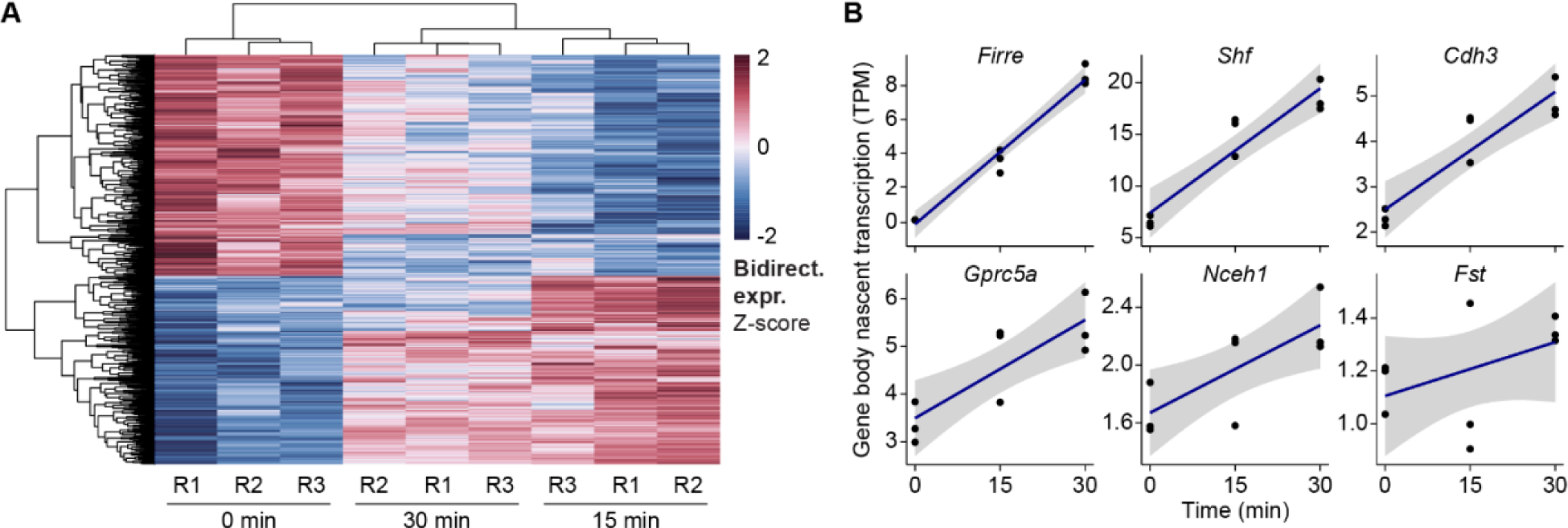
PRO-seq bidirectional and gene level changes. **(A)** Heatmap of the normalized counts +/- 300 bp from the bidirectional center for the 550 differentially expressed sites of bidirectional transcription. Counts are Z-scaled across rows. **(B)** Nascent transcription levels along the gene body across time. The trendline (blue) is fit with linear regression, and the standard error of the fit is shown in grey.

**Supplemental Figure S4:**
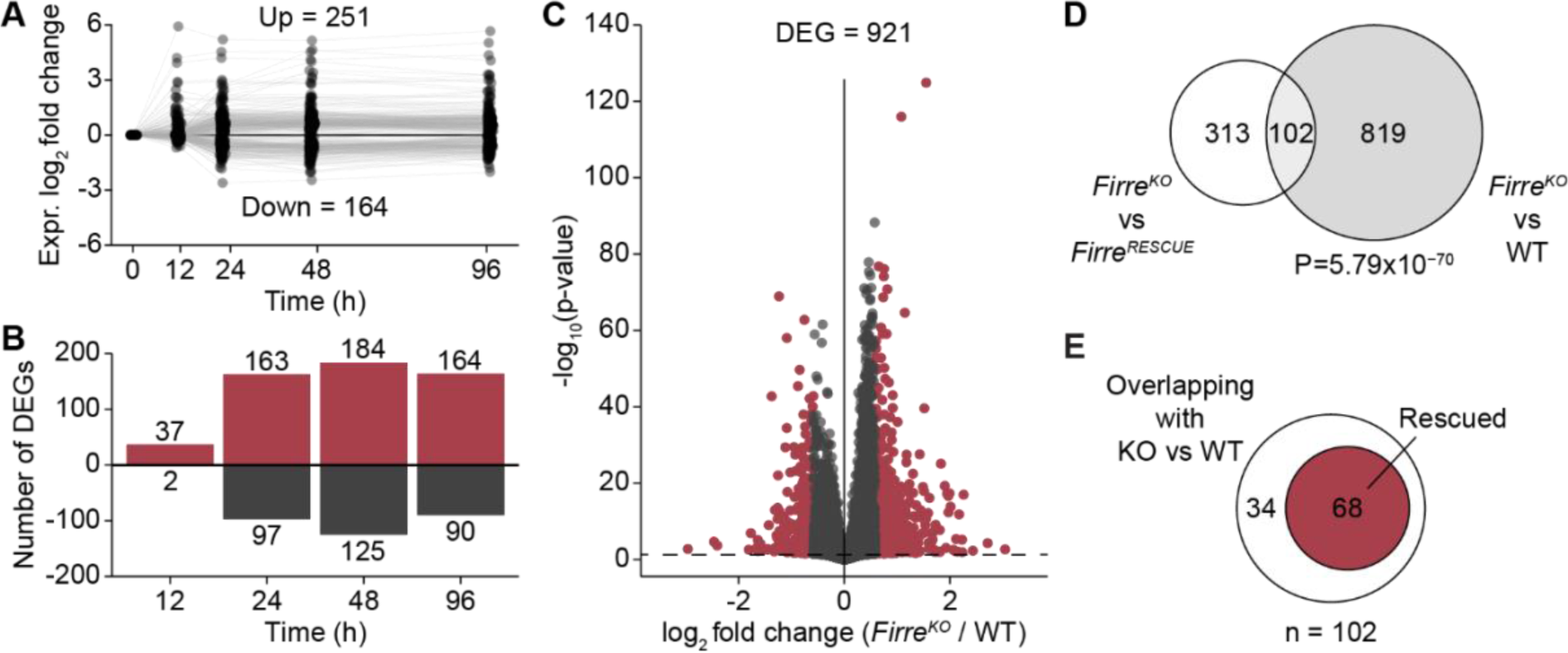
Late *Firre*-mediated gene regulation. **(A)** Time course plot of the shrunken log_2_ expression fold changes relative to the zero-time point of *Firre*-responsive genes in *Firre^RESCUE^*cells as determined by RNA-seq. **(B)** Bar plot showing the number of genes with a |fold change| > 1.5 at each time point in *Firre^RESCUE^*cells. Red and black bars indicate the number of upregulated and downregulated genes, respectively. **(C)** Volcano plot showing a total of 921 genes significantly differentially expressed in *Firre^KO^*relative to WT mESCs. *Firre* not shown: log_2_ fold change -6.9, -log_10_(p-value) = 196.6. **(D)** Venn diagram showing the overlap of significantly differentially expressed genes in *Firre^KO^* vs WT and *Firre^KO^* vs *Firre^RESCUE^* cells. **(E)** Venn diagram showing the number of rescued genes after induction of *Firre* compared to the total number of *Firre^KO^* vs WT overlapping differentially expressed genes.

## Methods

### Derivation and culture of genetically modified mESCs

mESC derivations were performed with the assistance of Harvard’s Genome Modification Facility. Briefly, 2-5 IU of pregnant mare serum gonadotropin (PMSG) and 2-5 IU of human chorionic gonadotropin (HCG) were administered 48 hours apart by intraperitoneal injection to female *Firre*- deficient (*Firre^-/-^*) mice. PMSG/HCG-treated mice were set up for timed matings (1-2 per cage) with either male mice deficient for *Firre* (*Firre^-/y^*) or with male mice that contained an inducible *Firre* transgene in the *Firre*-deficient background (*Firre^-/y^*; tg-*Firre*; *CAGs-rtTA3*) as previously described ^20^. Female mice were checked for copulation plugs the following morning. Approximately 72 hours later, blastocysts were flushed out of the uterine horns from female mice with successful copulation plugs. Blastocysts with a detected cavity were selected for culture and expansion. Individual clones were screened by PCR genotyping for *Firre* WT, KO, transgene, and *CAGs-rtTA3* alleles as described ^20^, and one independent cell line per genotype was derived. mESCs were maintained on 0.1% gelatin in KnockOut DMEM (Thermo Fisher, 10829018) supplemented with 12.5% FCS (MilliporeSigma, ES-009-B), 1X GlutaMAX supplement (Thermo Fisher, 35050061), 1X non-essential amino acids (Thermo Fisher, 11140050), 100 U/mL penicillin/streptomycin (Thermo Fisher, 15140122), 100 μM 2-mercaptoethanol (Thermo Fisher, 31350010), 1 μM PD0325901 (MilliporeSigma, PZ0162-5MG), 3 μM CHIR99021 (MilliporeSigma, SML1046-5MG), and 100 U/mL LIF (MilliporeSigma, ESG1107) at 37°C and 7.5% CO_2_. Cells were routinely tested for mycoplasma contamination by PCR.

Cells were treated with 1 ng/μL dox (MilliporeSigma, D9891-5G) the day after they were seeded and one hour after the medium was changed to reduce cell stress.

### RNA isolation, quantification, and quality control

Total RNA was extracted using TRIzol lysis reagent (Thermo Fisher) and purified using the RNeasy Plus Mini Kit (Qiagen) according to the manufacturer’s instructions. RNA concentration was measured with a Qubit fluorometer (Invitrogen), and RNA quality was determined using a Bioanalyzer or TapeStation (Agilent Technologies).

### RT-qPCR

Total RNA was reverse-transcribed using the SuperScript IV First-Strand Synthesis System with random hexamers (Thermo Fisher) according to the manufacturer’s instructions. RT-qPCR samples were prepared using TaqMan probes and the TaqMan Fast Advanced Master Mix (Thermo Fisher) according to the manufacturer’s instructions. The *Firre* transgene probe was designed against exons 1-2 and detects both the *Firre* transgene and endogenous *Firre*. Samples were run in technical triplicates on a Bio-Rad C1000 Touch Thermal Cycler. *C_t_*values were normalized against the internal controls *Gapdh* and *ActB*. Fold differences in expression levels were calculated according to the 2^−ΔΔ*Ct*^ method ^45^.

### RNA-seq

RNA-seq libraries were prepared from 1 μg of total RNA using the TruSeq RNA Library Prep Kit v2 (Illumina) with poly(A) selection according to the manufacturer’s instructions. Multiples of twelve randomized samples were each multiplexed, pooled, and run in one HiSeq lane.

### ATAC-seq

ATAC-seq libraries were prepared from 25,000 cells following the Omni-ATAC protocol ^46^ and using the Tagment DNA Enzyme and Buffer Kit (Illumina). Samples were pooled and run in one HiSeq lane.

### PRO-seq

Nuclei were purified by washing cells twice with cold PBS and incubating them in ice-cold swelling buffer (10 mM Tris-HCl pH 7.5, 2 mM MgCl_2_, 3 mM CaCl_2_) on ice for 15 min. Cells were collected in 20 mL swelling buffer using a cell scraper and centrifuged at 1000 x g for 10 min at 4°C. Cell pellets were resuspended in 1 mL of lysis buffer (10 mM Tris HCl pH 7.5, 2 mM MgCl_2,_ 3 mM CaCl_2_, 0.5% IGEPAL, 10% glycerol, RNase Inhibitor) using a wide-orifice pipette tip. An additional 9 mL of lysis buffer was added, and the mixture was centrifuged at 600 x g for 5 min at 4°C. This resuspension step was repeated once. The nuclei pellets were then resuspended in 1 mL of freezing buffer (50 mM Tris HCl pH 8.3, 5 mM MgCl_2_, 40% glycerol, 0.1 mM EDTA pH 8.0) and transferred to low-bind microcentrifuge tubes. After centrifugation at 600 x g at 4°C for 5 min, nuclei pellets were resuspended in 500 μL of freezing buffer and centrifuged at 600 x g for 5 min at 4°C. Nuclei pellets were finally resuspended in 110 μL of freezing buffer and flash-frozen in liquid nitrogen to be stored overnight at -80°C.

The nuclear run-on experiments for PRO-seq were performed in triplicate as described in ^47^ with the use of a mixture of rNTPs and Biotin-11-CTP (PerkinElmer), and a range of 10-20 million isolated nuclei per sample. Briefly, after the nuclear transcription run-on reaction, total RNA was extracted with TRIzol LS reagent (Thermo Fisher) and then fragmented by base hydrolysis. Biotinylated fragmented nascent transcripts were enriched a total of three times using streptavidin Dynabeads (Invitrogen) and TRIzol LS reagent. The VRA3 RNA adaptor was ligated to the 3’ end of nascent RNA following the first biotin enrichment. After the second biotin enrichment, the 5 ’ends of the RNA were enzymatically modified with pyrophosphohydrolase and polynucleotide kinase and then ligated to the VRA5 RNA adaptor. Following the third biotin enrichment, a reverse transcription reaction was performed, and the resulting adaptor-ligated libraries were purified using AMPure XP beads (Beckman Coulter). The libraries were amplified with eleven cycles of PCR, purified with AMPure beads, pooled, and run on the NovaSeq platform.

### smRNA FISH

Oligonucleotides tiling *Firre* exons (including its RRDs) and *Shf* exons were designed with the Stellaris RNA FISH probe designer (LGC Biosearch Technologies, version 4.2), labeled with Quasar 570 or 670, and produced by LGC Biosearch Technologies. Probe sequences are provided in Supplemental Table S1.

Cells were seeded on cover glasses coated with 0.1% gelatin. Cover glasses were washed twice with PBS, fixed in 3.7% formaldehyde in PBS for 10 min at room temperature, and washed again twice with PBS. Cover glasses were incubated in 70% ethanol at 4°C for at least one hour and then washed with 1 mL of wash buffer A (LGC Biosearch Technologies) at room temperature for 5 min. Cells were hybridized with 100 μL of hybridization buffer (LGC Biosearch Technologies) containing the FISH probes at a 1:100 dilution in a humid chamber at 37 °C overnight up to 16 hours. The next day, cells were washed with 1 mL of wash buffer A at 37 °C for 30 min and stained with wash buffer A containing 10 μg/mL Hoechst 33342 (Thermo Fisher) at 37 °C for 30 min. Cover glasses were washed with 1 mL of wash buffer B (LGC Biosearch Technologies) at room temperature for 5 min, mounted with ProLong Gold (Thermo Fisher) on a glass slide, and left to curate at 4°C overnight before proceeding to image acquisition.

Image acquisition was performed using a DeltaVision Elite widefield microscope with an Olympus UPlanSApo 100×/1.40-numerical aperture oil objective lens and a PCO Edge sCMOS camera. Z-stacks of 200-nm step size capturing the entire cell colony were acquired. Images were deconvolved with the built-in DeltaVision SoftWoRx Imaging software and maximum intensity projections were created using ImageJ/Fiji. FISH spots were quantified manually using ImageJ/Fiji.

## Data analysis

### RNA-seq analysis

Quality control, read mapping, and quantification was performed using nf-core/rnaseq v1.4.2 ^48^. Reads were mapped to mm10 and quantified with Salmon v1.5.2 ^49^ using the Gencode M25 annotation.

Static *Firre^KO^*vs WT analysis: Significantly differentially expressed genes were calculated using DESeq2 ^50^ with an adjusted p-value cutoff of 0.05 and a fold change cutoff of 1.5 on the shrunken log_2_ fold change values from DESeq2.

Dox control analysis: Differential expression was calculated using DEseq2 and the Wald test to compare each time point back to the zero-time point. The rescue and overexpression dox control lines and the corresponding short and long-time courses were analyzed separately. Cutoffs to call differentially expressed genes were made using an adjusted p-value cutoff of 0.05 and whether a gene achieved a fold change greater than 1.5 at any time point (calculated using the shrunken log_2_ fold change).

*Firre* induction time course analysis: Differential expression was calculated with DEseq2 using the likelihood ratio test to compare a full model which contains the *Firre* induction/time point interaction term to a reduced model lacking that term. Calling differentially expressed genes proceeded in two steps: 1. A cutoff was made on the adjusted p-values from the likelihood ratio test (p < 0.05), filtering to genes that changed significantly in time as compared to the dox control. 2. A fold change cutoff was made by comparing the expression through time back to the zero - time point (without any comparison to the dox control line). A gene was called differentially expressed (responding to *Firre*) if it achieved a fold change greater than 1.5 in any time point as measured by the shrunken log_2_ fold change from DEseq2.

The combined model using all four cell lines was analyzed in the same manner with the same thresholding strategy, but the full and reduced models contained an additional indicator variable for the *Firre* genotype (WT or KO).

The log_2_ fold changes shown in the heatmaps are the shrunken log_2_ fold changes reported by DEseq2.

Identification of rescued genes: A gene was considered rescued, if: 1. It was significantly differentially expressed in the static *Firre^KO^*vs WT comparison and in the *Firre^RESCUE^* induction experiment, and 2. If the fold change was reciprocal to within 20% of the fold change value in the *Firre^KO^* vs WT.

To determine whether any early Firre responding genes may result from rTTA activation, publicly available RNA-seq count data were retrieved from the Gene Expression Omnibus (GEO) database for the accession number GSE118898. Specifically, raw RNA-seq count data for the following samples were extracted: (HV_G1_SunTag_Nanog_A1_pLif_mDox_1, HV_G5_SunTag_Nanog_E1_pLif_mDox_1, HV_G7_SunTag_Nanog_E1_pLif_mDox_2, HV_G4_SunTag_Nanog_A1_pLif_pDox_2, HV_G6_SunTag_Nanog_E1_pLif_pDox_1).

These samples represent conditions with (+Dox) and without doxycycline (-Dox) treatment in a mouse embryonic stem cell line (E14Tg2a) containing a dox-inducible Nanog transgene. Differential expression analysis between the pDox and mDox conditions was performed using the DESeq2 R package.

### ATAC-seq analysis

Quality control, read mapping, and peak calling were carried out using nf-core/atacseq v1.2.0. Reads were mapped to mm10 and the mm10 v2 blacklist from ENCODE was used. Within the nf-core/atacseq pipeline MACS2 v2.2.7.1 ^51,52^ was used to call peaks in each experiment separately. Consensus peaks were called using the peak regions present in any single experiment. Reads within peaks were calculated using featureCounts. Differentially accessible peaks were calculated using DEseq2 to compare the read counts in the samples in which *Firre* was present (>0 time point in the *Firre^RESCUE^* and *Firre^OE^*lines) and the samples in which *Firre* was absent (WT and *Firre^KO-CTL^*lines, and zero-time points). A cutoff was made using the adjusted p-value (p<0.05).

### PRO-seq analysis

Quality control and read mapping were performed using the Nascent-Flow Nextflow pipeline (https://github.com/Dowell-Lab/Nascent-Flow). In that pipeline, reads were mapped to mm10 using Hisat2 with the flags --very-sensitive and --no-spliced-alignment. Bidirectionals were called with Tfit using the Bidirectional-Flow pipeline with the default parameters (https://github.com/Dowell-Lab/Nascent-Flow). muMerge v1.1.0 (https://github.com/Dowell-Lab/mumerge) was used to merge bidirectional nascent transcript calls across samples. For quantification, a uniform 600 bp region from the center of the muMerge identified bidirectional was used. Counts in each sample were called using Rsubread’s (v2.0.1) featureCounts over these regions. To filter to transcripts that produced reads on both transcripts, featureCounts was used to generate counts over the positive and negative strand aligned reads by using separate ’saf’ files over these 600 bp regions with + and - strand regions. Bidirectionals were then filtered to those that had >=5 reads on each strand and a strand bias ratio of <1. To identify differentially expressed nascent transcripts over the time course, DESeq2 (v1.26.0) was used to calculate a likelihood ratio test between the full model (∼timepoint) vs a reduced model (∼1). Further, to evaluate the magnitude of expression changes, DESeq2 was run using the Wald test to compare the 15- and 30-min time points to the zero-time point. A transcript was considered significant if the LRT test padj < 0.05 and achieved |fold change| > 1.5 in either time point. The metaplot of PRO-seq signal over the 550 differential bidirectional regions was generated by summing the signal over the positive and negative strands separately over all replicates. To detect changes in nascent transcription along the gene body, Rsubread’s (v2.0.1) featureCounts was used to quantify counts over the gene body lacking the 5’-most 500 bp so as not to include the promoter bidirectional. Differential expression of nascent transcription in the gene body was performed with DESeq2 (v1.26.0) using a likelihood ratio test over the time points with the full model (∼timepoint) and reduced model (∼1).

## Data access

All sequencing data generated in this study have been submitted to the NCBI Gene Expression Omnibus (GEO; https://www.ncbi.nlm.nih.gov/geo/) under accession number GSE202406. All code to reproduce the analyses is available at: github.com/msmallegan/firre_timecourse.

## Acknowledgements

We would like to thank Gabrijela Dumbović and Aaron Nguyen for their help with smRNA FISH, Nydia Chang for assistance in deriving the mESCs, and Taeyoung Hwang for his initial guidance in analyzing the *Firre* time course data. We are grateful to Roy Parker and Carolyn Decker for access to, and training on, the DeltaVision Elite microscope. We acknowledge the support of Theresa Nahreini and Nicole Kethley at the Biochemistry Cell Culture Facility, and Amber Scott at the BioFrontiers Sequencing Core. C.M. was supported by the Deutsche Forschungsgemeinschaft (DFG, MU 4462/1-1). J.R. is HHMI Faculty Scholar and the work was supported by NIH/NIGMS grant P01 GM099117.

